# What Does Large-scale Electrodermal Sensing Reveal?

**DOI:** 10.1101/2024.02.22.581472

**Authors:** Daniel McDuff, Seamus Thomson, Samy Abdel-Ghaffar, Isaac R. Galatzer-Levy, Ming-Zher Poh, Jake Sunshine, Andrew Barakat, Conor Heneghan, Lindsey Sunden

**Affiliations:** Google Inc., USA

## Abstract

Electrodermal activity (EDA) is a physiological measure that is used to index sympathetic arousal in response to stressors and other perturbations. However, EDA is underutilized in real-world, population-level research and clinical practice because of a paucity of remote measurement capabilities on commodity devices. The current study examined the capabilities of continuous remote measurement of EDA at scale to quantify physiological changes in the context of diurnal and circadian rhythms, demographic differences, high arousal contexts such as public holidays and high arousal moments (e.g., the Super Bowl). We first demonstrated the accuracy of a novel EDA sensor developed for the Fitbit Sense 2 wearable device within a controlled, arousal-inducing experiment. We then retrospectively analyzed 10 million hours of continuous EDA data collected from over 16,000 people. We show that continuously sampled *in-situ* EDA from Sense 2 has similar population-level diurnal patterns as those established from more tightly controlled in-lab experiments. Following this, variation of SCL across day of the week and season are presented. Finally, EDA dynamics occurring in response to commonly held stressful or exciting events such as Thanksgiving and the Super Bowl are described which we interpret as a natural experiment eliciting autonomic arousal.

## Introduction

Electrodermal activity (EDA) has long been utilized in psychological research as a physiological index of autonomic arousal and stress [Boucsein, 2012]. Chronic stress is known to be associated with many negative health outcomes, some of which include: heart disease, hypertension, diabetes and poor sleep. Quantitative measurement of physiological stress and arousal thus has applications within basic and clinical psychological research. The measurement of EDA [Johnson and Lubin, 1966] was standardized by the Society for Psychophysiological Research in the late 1960s [Brown, 1967]. EDA is measured by placing electrodes on the skin and recording the electrical potential between them. This electrical potential changes in response to changes in sweat gland activity. When a person is stressed or aroused, there is an increase in activity of the sympathetic branch of the autonomic nervous system that results in the production of sweat, which causes an increase in the electrical potential of the skin. By measuring this change in electrical potentials across the skin, EDA can be used to track sympathetic nervous system arousal over time, or to compare arousal levels between different people or groups. EDA has been evaluated, to a limited extent, over 24-hour periods [Vieluf et al., 2021, Posada-Quintero and Chon, 2020, Klimek et al., 2023]. Yet, despite its value as an index across psychological science, it is under-studied and under-utilized within long-duration, in-the-wild and population level research because it was unavailable through remotely connected, commodity smart devices.

### Real-World Measurement of EDA

Within laboratory settings, researchers measuring EDA typically use gel electrodes to reduce the effect of skin-electrode impedance, and hence reduce measurement error; however, this cumbersome interface limits real-world applications at scale. The emergence of dry electrode technology has facilitated EDA measurement without the need for the application of gel. Rather, dry electrodes are directly affixed to the skin and rely on the wearable device to hold the sensor in place, allowing for real-world continuous measurement of EDA [Poh et al., 2010, Garbarino et al., 2014]. The Affectiva Q sensor^1^ and Empatica E4^2^ pioneered wrist-worn dry electrode measurement and these sensors have been employed in psychophysiological studies. However, many of these studies focus on specific populations and/or small cohorts [Regalia et al., 2019, Halimeh et al., 2022, Ahmadi et al., 2022, Vieluf et al., 2021] making it difficult to draw robust conclusions about typical variations in everyday life. The availability of EDA sensors on popular smartwatches ^3^, and smartrings^4^ now enables population study of EDA for health and research applications.

### Why is measuring sympathetic arousal significant?

EDA is commonly utilized in laboratory and clinical settings to quantify psycho-physiological arousal, and has been found to vary with sleep and circadian rhythms, to change in response to acute psychological stress and pain, and to differ significantly across neurological and psychiatric conditions [Tronstad et al., 2022, Vieluf et al., 2021]. Specifically, tonic changes in the EDA signal - represented by the Skin Conductance Level (SCL) - have been reported as a useful feature for evaluating sympathetic arousal [Kreibig, 2010, Jacobs et al., 1994]. Measuring EDA continuously and reliably in real-world settings has the potential to advance knowledge of normal and pathological stress and arousal patterns and, importantly, their associations with downstream health outcomes. As one remarkable example, Carroll et al. (2002) observed that the risk of admission to hospital for acute myocardial infarction increased by 25 percent the day after England lost to Argentina in a penalty shoot-out in the 1998 FIFA World Cup. Newson et al. (2020) measured salivary cortisol changes during the 2014 FIFA World Cup and found an effect of match outcome on cortisol, where a loss was associated with higher cortisol concentrations. However, evaluating large-scale *in-situ* stress exposure via salivary sampling is challenging due to the logistical challenges associated with the collection of bodily samples for chemical analysis. EDA measurement would be a practical, scalable alternative. Indeed, in emergency medical settings in the immediate aftermath of accidents and injuries, EDA been shown to be predictive of the development of psychiatric sequelae including PTSD [Hinrichs et al., 2017].

Real-world measurement of EDA presents distinct challenges, as perspiration is not specific to stress or sympathetic arousal. Specifically, ambient temperature and activity both impact individual differences in SCL, a result of the body’s thermoregulatory capabilities. These factors must be measured and accounted for via other sensor modalities in order to reliably index psychophysiological responses, as opposed to confounding sources of SCL variability. Given these experimental and technological challenges, limited observational evidence exists regarding how EDA varies in everyday life. Pioneering studies of continuously measuring EDA do show a pattern of change over a 24-hour period, with the highest SCL observed at night [Vieluf et al., 2021]. During daytime hours, SCL is found to be higher at mid-day and the early afternoon/evening than in the morning [Hot et al., 1999]. However, these analyses do not all control for confounding variables (e.g. activity patterns) nor reveal how these patterns vary based on their covariance with other variables of interest known to affect arousal (e.g. day of the week, season, BMI, age, gender and other physiological measures).

In this work, we first demonstrate the accuracy of a novel EDA sensor developed for the Fitbit Sense 2 wearable device within a controlled, arousal-inducing experiment. Next we show that continuously sampled *in-situ* EDA from Sense 2 has similar population-level diurnal patterns as those established from more tightly controlled in-lab experiments [Hot et al., 1999, 2005]. Following this, variation of SCL across day of the week and season is presented. Finally, we present population-level EDA dynamics in response to commonly held stressful or exciting events such as Thanksgiving and the Super Bowl which we interpret as a natural experiment eliciting autonomic arousal.

## Results

### Validation During a Controlled Laboratory Stress Induction Experiment

To validate the measurement of SCL from the dorsal wrist we performed a Trier Social Stress Test (TSST) [Kirschbaum et al., 1993] experiment. Individuals (N=45; men=20, age: 38.2 (*σ*=8.8) years, min/max age: 24/60) completed the TSST protocol remotely while wearing a Fitbit Sense 2 (see Methods for details). This remote protocol has previously been validated and was a necessary constraint as a result of the COVID-19 pandemic [Eagle et al., 2021]. Full details about the experiment can be found in the Appendix. The EDA signal was acquired using a Fitbit Sense 2 during the baseline and stressor conditions of the test. A two-sided Mann-Whitney U test was performed to evaluate whether SCL differed by condition. Consistent with expectations, the results indicated that SCL was significantly greater during stress tasks than during the baseline period (*p* = .004 based on a Mann-Whitney U Test). The mean SCL during the baseline period was 1.47*μ*Siemens and during the stress period was 2.17*μ*Siemens. These results provide confidence that tonic SCL measured from the dorsal wrist increases during a known and validated stressor.

### Population Assessment of Electrodermal Activity

Next, we moved to analyze data from our *in-situ* population (N=15,349 users). To summarize the population, the users were a majority women (65.5%) and skewed younger than the U.S. population (see Table 1); however, there were a large number of men (5,300) and older adults (2,462 people 65 year or older). The body-mass index (BMI) distribution was skewed slightly higher than the Center for Disease Control (CDC) population estimates. BMI statistics are based on user’s self-report and may have some measurement error.

**Table 1:** A summary of the demographics of the *in-situ* users in our study. We compared these to population distributions.

| Category | No of Individuals | Percentage | U.S. National Census Data (2021) |
| --- | --- | --- | --- |
| **Gender** |  |  |  |
| Female | 10,049 | 65.5% | 50.4% |
| Male | 5,300 | 34.5% | 49.6% |
| **Age** |  |  |  |
| 18-65 | 12,966 | 84.5% | 77.9% |
| 65+ | 2,383 | 15.4% | 22.1% |
| **BMI** |  |  |  |
| Under 25 (Healthy) | 3,260 | 20.3% | 32.5%<sup>†</sup> |
| 25-30 (Overweight) | 4,957 | 30.9% | 35.4%<sup>†</sup> |
| 30+ (Obese) | 6,169 | 40.2% | 32.1%<sup>†</sup> |
| Unknown | 963 | 6.3% |  |

<sup>†</sup> https://www.cdc.gov/nchs/nhis/2022nhis.htm

We examined the distribution of minutely SCL levels across the entire population (mean = 4.23 *μ*S; median = 3.34 *μ*S; Percentiles = 5th: 0.96 *μ*S, 10th: 1.27 *μ*S, 90th: 8.44 *μ*S, 95th: 10.66 *μ*S). These statistics reflect a distribution with slightly higher mean SCL than previous ambulatory studies [Vieluf et al., 2021]; which may be explained by this being a slightly more active population (as is the typical user base for fitness trackers). A plot of the population mean daily SCL (see Fig. 2) revealed that warmer months appear to demonstrate elevated SCL and higher SCL variance compared to colder months. Zooming into the time axis we see rhythmic differences by weekday and weekend where weekends appear to produce higher mean SCL scores. Zooming in further still reveals distinct repetition each day, or diurnal patterns.

**Figure 1:**
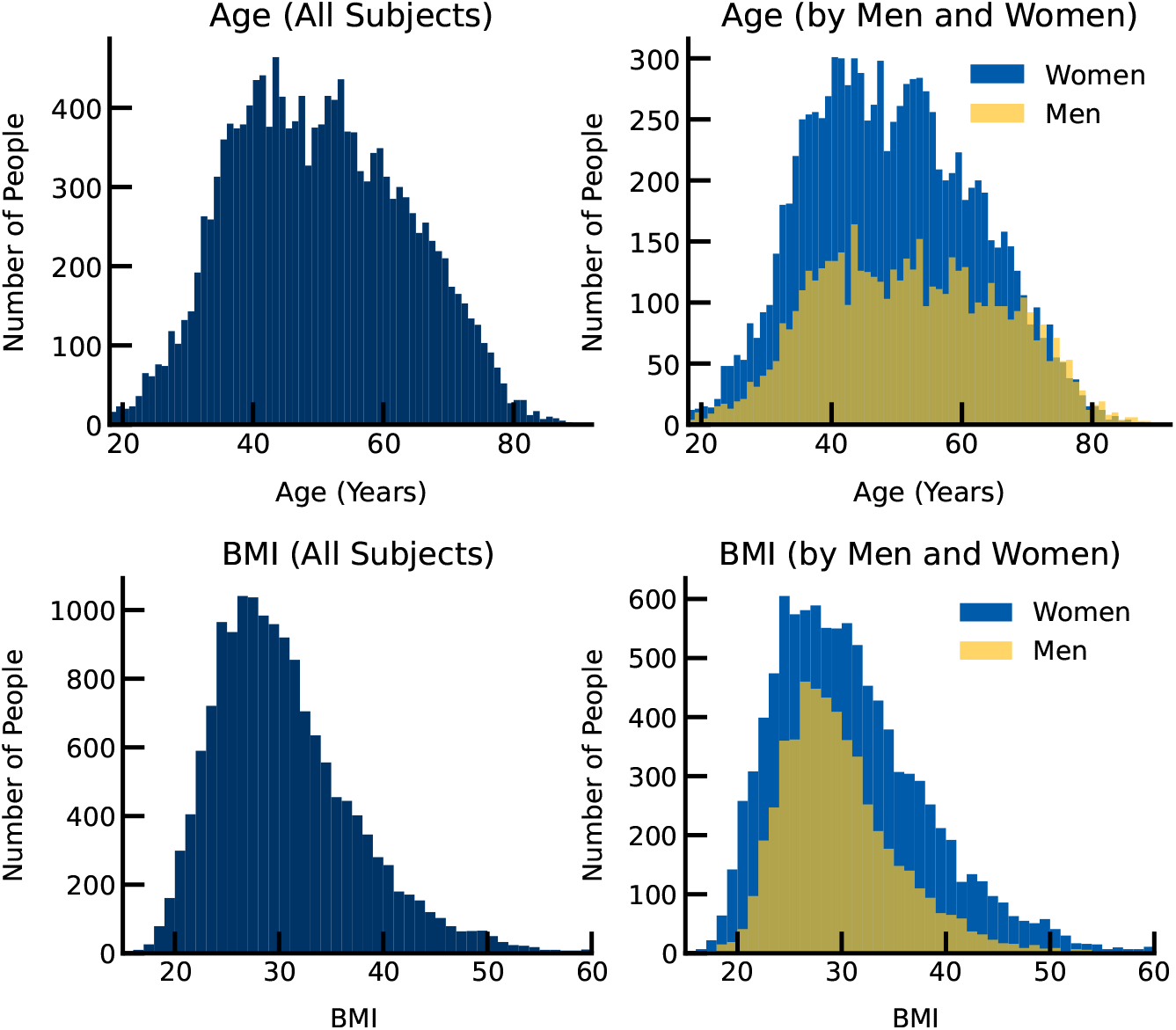
Distributions of the age and BMI of the individuals in our sample, overall (left) and by gender (right).

**Figure 2:**
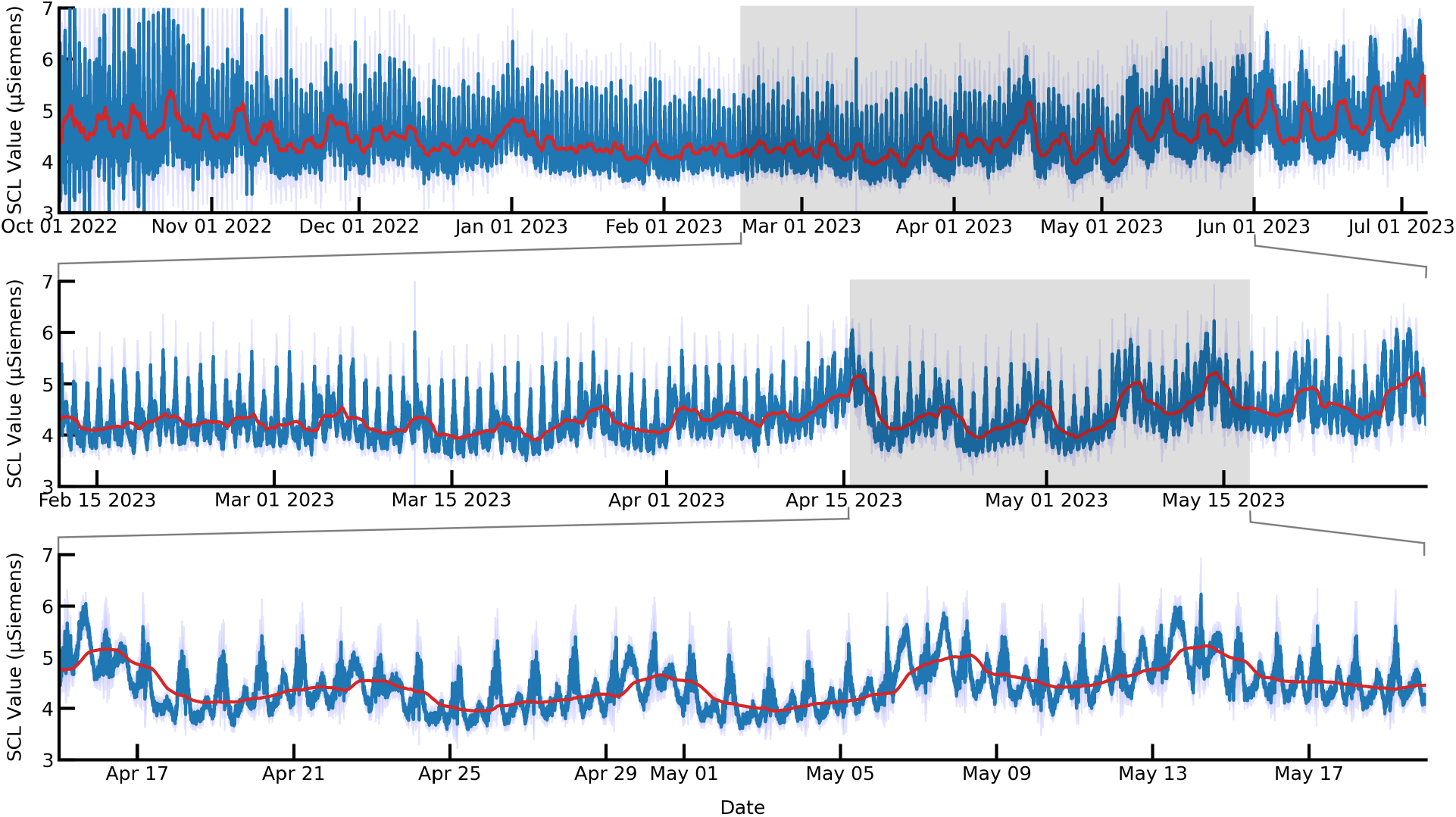
Population SCL mean from October 1st 2022 to July 1st 2023. One-day moving mean average shown in red. SCL levels are higher and show greater variance in warmer months. Weekends have higher SCL than weekdays and nighttime hours (8pm - 8am) have higher SCL than daytime hours (8am - 8pm). Shaded areas represent 95% confidence based on standard error.

A multiple linear regression was applied to assess the relationship of known covariates of SCL including demographics (Gender, Age, BMI), other physiological processes related to sympathetic arousal (skin temperature, heart rate, heart rate variability), behavior (steps, sleep duration), and time of day (dummy coded at three levels representing morning (6am - 12pm), afternoon (12pm-6pm), and night (6pm -12am), time of week (dummy coded comparing weekdays to weekend), and season (dummy coded comparing spring, summer and fall to winter) along with two-way interactions between demographics (age × gender) and time (time of day × time and day of week). Based on evidence that the SCL values were not normally distributed and had a positive skew, we performed a Box-Cox transformation after which normality tests were satisfied (Shapiro-Wilks: *p* = 0.210, histograms and QQ-Plots can be found in the supplemental materials). We also tested the distributions of the model residuals and confirmed that they satisfied a normal distribution (two-sided Jarque-Bera chi^2^ test, *p*=0.209).

Consistent with prior literature, results (Table 2, Model V) demonstrated main effects for demographics with BMI, and female gender demonstrating positive linear relationships with hourly tonic SCL. A significant interaction was detected between gender and age. Mean physiological data across the day also demonstrated linear relationships with mean SCL whereby skin temperature, heart rate, and step count all demonstrated positive associations with SCL while HRV, which decreases when sympathetic arousal is elevated, demonstrated a negative linear relationship with EDA (Table 1). These results support prior findings on correlations between skin conductance response (SCR) amplitude and HR [Ahmadi et al., 2022]. Time in bed was also negatively correlated with SCL.

**Table 2:** Linear model fit via ordinary least squares regression. Component (demographics, physiology and time/day of week) models indicate that demographics and physiology explain a higher proportion of the variance in hourly tonic SCL.

| Predictor Variable | (I)<br>Demograph | (II)<br>Phys/Behav | (III)<br>Time/Day | (IV)<br>Full (No Ints.) | (V)<br>Full (Ints.) |
| --- | --- | --- | --- | --- | --- |
| *Intercept* | 1.197*** | 1.107*** | 1.039*** | 1.270*** | 1.219*** |
| **Demographics** |  |  |  |  |  |
| *BMI* | 0.133*** |  |  | 0.141*** | 0.142*** |
| *Age* | -0.012*** |  |  | -0.001 | 0.036*** |
| *Gender (Male)* | -0.243*** |  |  | -0.259*** | -0.256*** |
| *Age\*Gender* |  |  |  |  | -0.090*** |
| **Physiology & Behavior** |  |  |  |  | ... |
| *Skin Temp* |  | 0.156*** |  | 0.180*** | 0.179*** |
| *Heart Rate* |  | 0.095*** |  | 0.063*** | 0.063*** |
| *HRV SDNN* |  | -0.059*** |  | -0.035*** | -0.034*** |
| *Step Count* |  | 0.116*** |  | 0.150*** | 0.149*** |
| *Time in Bed* |  | -0.028*** |  | -0.028*** | -0.027*** |
| **Time** |  |  |  |  | ... |
| *6am-12pm* |  |  | 0.084*** | 0.006** | 0.074*** |
| *12pm-6pm* |  |  | 0.135*** | -0.068*** | 0.026 |
| *6pm-12am* |  |  | 0.062*** | -0.070*** | -0.036*** |
| *Day of Week (Mon-Fri)* |  |  | -0.083*** | -0.068*** | 0.007** |
| *Spring* |  |  | 0.036*** | 0.001 | 0.002 |
| *Summer* |  |  | 0.142*** | 0.058*** | 0.059*** |
| *Fall* |  |  | 0.065*** | 0.056*** | 0.055*** |
| *6am-12pm\*Weekday* |  |  |  |  | -0.096*** |
| *12pm-6pm\*Weekday* |  |  |  |  | -0.132*** |
| *6pm-12am\*Weekday* |  |  |  |  | -0.048*** |
| Marginal R<sup>2</sup> | 0.037 | 0.061 | 0.007 | 0.102 | 0.105 |
| Number of Observations | 433,154 | 433,154 | 433,154 | 433,154 | 433,154 |

### Temporal Patterns

#### Diurnal Patterns

Given the significance of diurnal and circadian rhythms in biology, we directly analysed diurnal rhythms across 24-hour periods and the differences between those patterns on weekdays and weekends. To support the analysis of diurnal variability, patterns for SCL were compared to those for other physiological measures, namely: HR, HRV, steps, and skin temperature.

The diurnal rhythm was calculated using the 24 hour day/night cycle based on the local time of each user. Circadian patterns were calculated by adjusting for the wake time of the user each day and averaging subject’s observations over time into a single trajectory (see Fig. 3). As is common practice, a cosine function was fit with ordinary least squares to each measure of physiology (SCL, HRV, HR, Steps, skin temperature). Results demonstrated distinct trajectories for each metric with changes during sleep/wake hours, and across the day. When comparing weekdays to weekends, SCL demonstrated a distinct elevation during the day. Based on the results of our previous model and the diurnal plots the changes in SCL are not consistent or attributable in a large part to variability in skin temp or movement, or cardiovascular output. The pattern for SCL had a nadir near 10pm local time and a zenith near 6am local time. But notably, whereas other metrics were best fit with a cosine of wavelength equal to one day, SCL was best fit with a half-day wavelength, indicating two maxima across the course of the day. A secondary nadir was observed at 10am and a zenith at 6pm. Furthermore, these patterns deviated significantly from weekdays compared to weekends, more so than those for other physiological signals (see Fig. 3ii and iii). These results suggest that differences in activity and environment between the weekdays and weekends disproportionately effect mean SCL. Hypotheses about the reasons for these large differences are discussed later.

**Figure 3:**
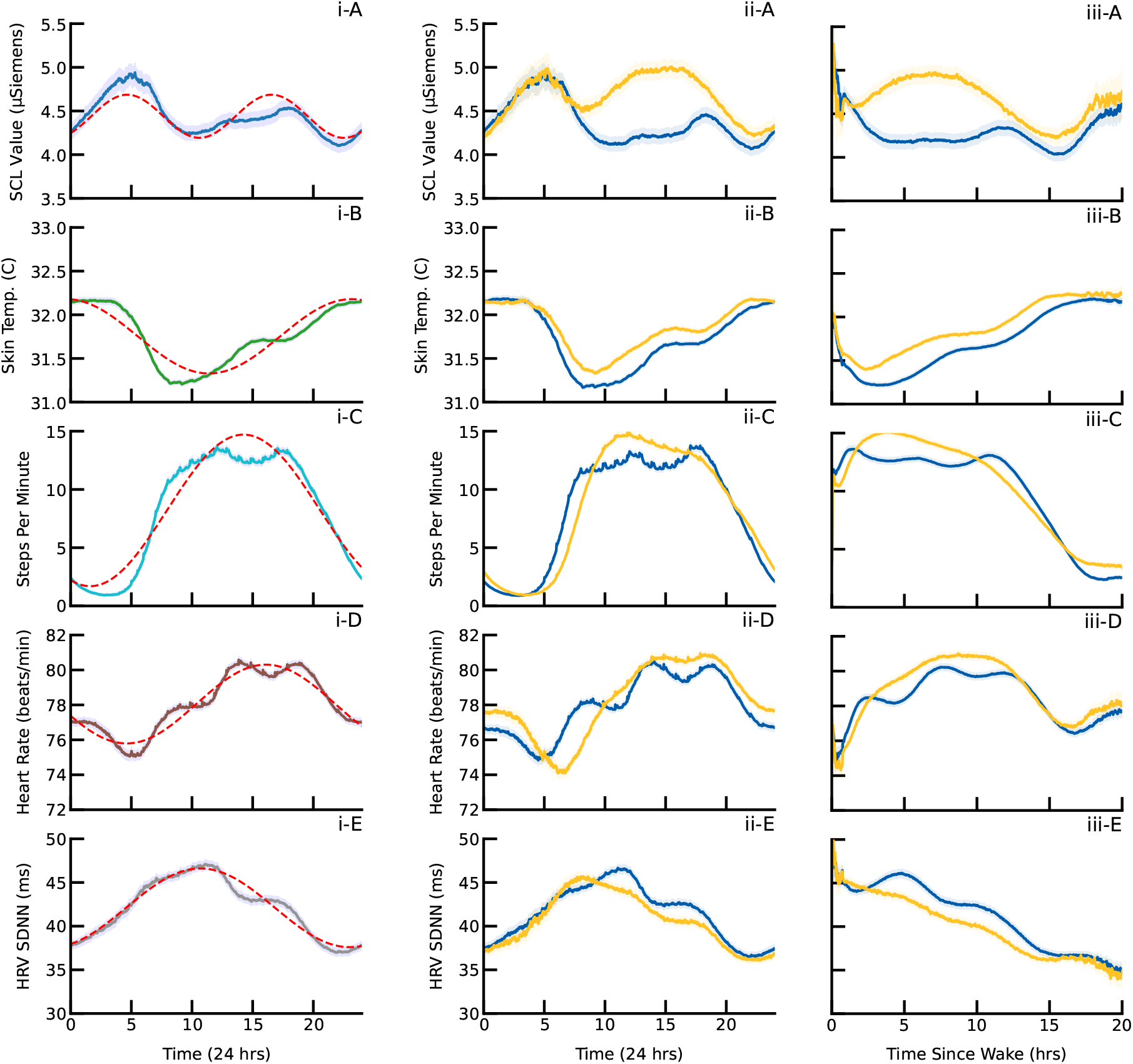
Mean (i) 24 hour period across all days (red dashed line - cosine best fit), (ii) 24 hour period by weekdays (blue) and weekends (yellow), and (iii) time since wake patterns. (A) Tonic SCL, (B) skin temperature, (C) Steps, (D) HR, and (E) HRV SDNN for Monday-Friday (blue) and Saturday-Sunday (yellow). Shaded areas represent 95% confidence based on standard error.

#### Gender and Age

To better understand the interaction between age and gender, diurnal trajectories were plotted and statistically compared between gender and age (<=35 or => 50 y.o.). The age threshold was chosen to highlight differences between young adults and middle age to older adults. The threshold of 50 y.o. (median age at menopause is approximately 51.5 years [Bromberger et al., 1997]) was chosen because of evidence that hot flashes are significantly associated with the onset of menopause [Avis et al., 2003], and that these in turn influence SCL [Thurston et al., 2009]. Our model indicated that overall women have higher SCL than men, and this effect is driven by large increases starting in the fifth decade of life (see Fig. 4).

**Figure 4:**
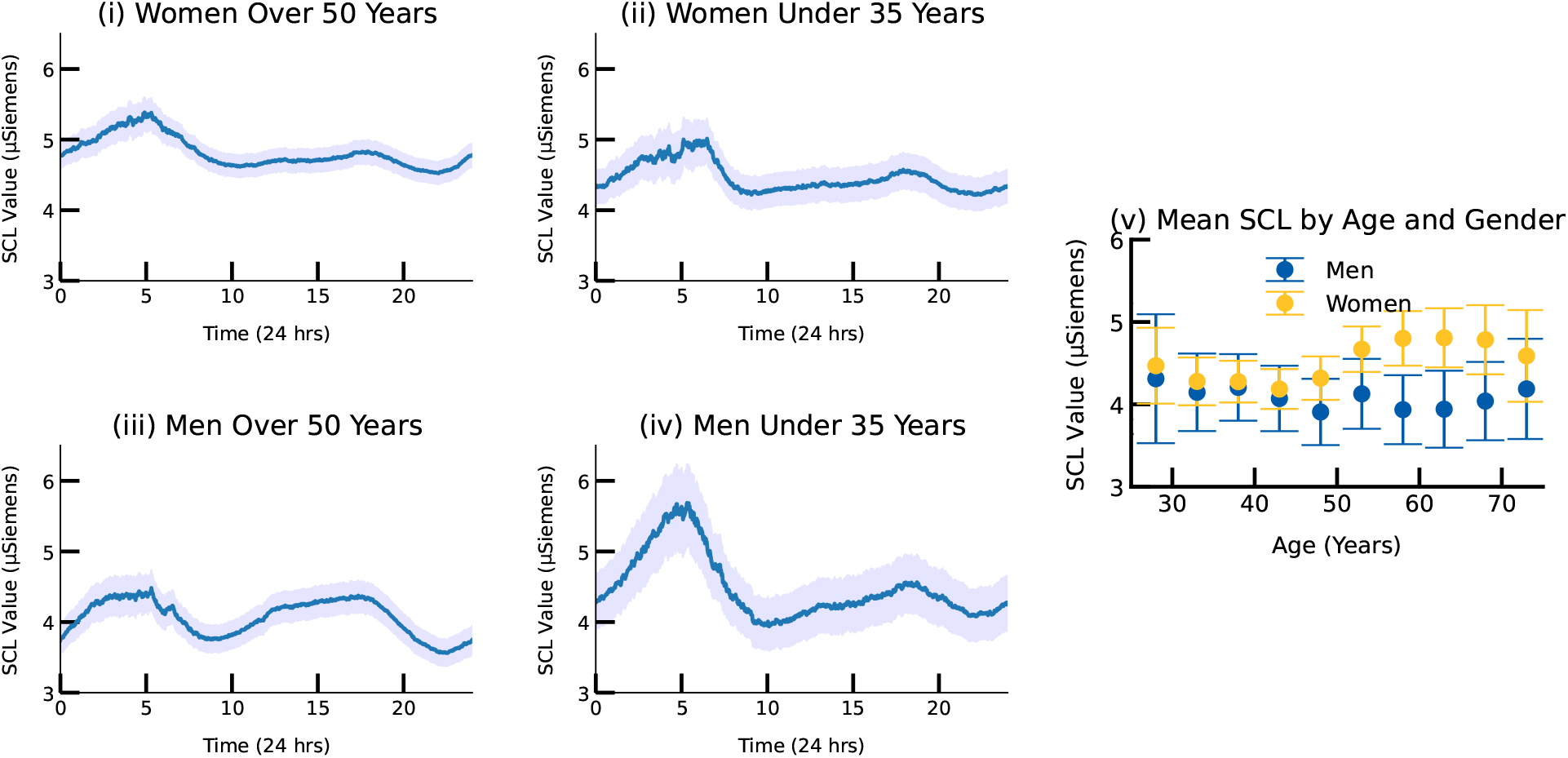
Mean diurnal patterns by 24 hour period in tonic SCL for (i-ii) women and (iii-iv) men and those over 50 years (i and iii) and under 35 years (ii and iv). (v) Mean SCL by age and gender. Mean SCL increased for those self-identified as women over the age of 50 years. Shaded areas and error bars represent 95% confidence based on standard error.

### High Arousal Events

To illustrate the effect of stressful events on SCL, we identified common, time-limited stressful life events that are shared across a population. Thanksgiving Day and Super Bowl LVII are two such events analyzed here.

#### Thanksgiving

To analyse the effects of Thanksgiving Day, subject data was mean averaged across the 24 hour period of Thanksgiving Day 2022 (November 24^th^), and statistically compared to the same time range collected on other Thursdays in November and December 2022 (see Fig. 5A). The largest differences between typical Thursday diurnal SCL were between 11am-2pm local time. SCL was significantly higher on Thanksgiving Day compared to an average Thursday. A two-sided Mann-Whitney U test was performed to evaluate whether SCL differed between Thanksgiving and other fall Thursdays and the results indicated that Thanksgiving had significantly greater SCL than other fall Thursdays (z = [219,418], *p* < [.001]). As Thanksgiving day is a national holiday, we also compared mean SCL to that on Saturdays in November and December, again SCL was significantly higher on Thanksgiving Day (z = [324,591], *p* < [.001]).

**Figure 5:**
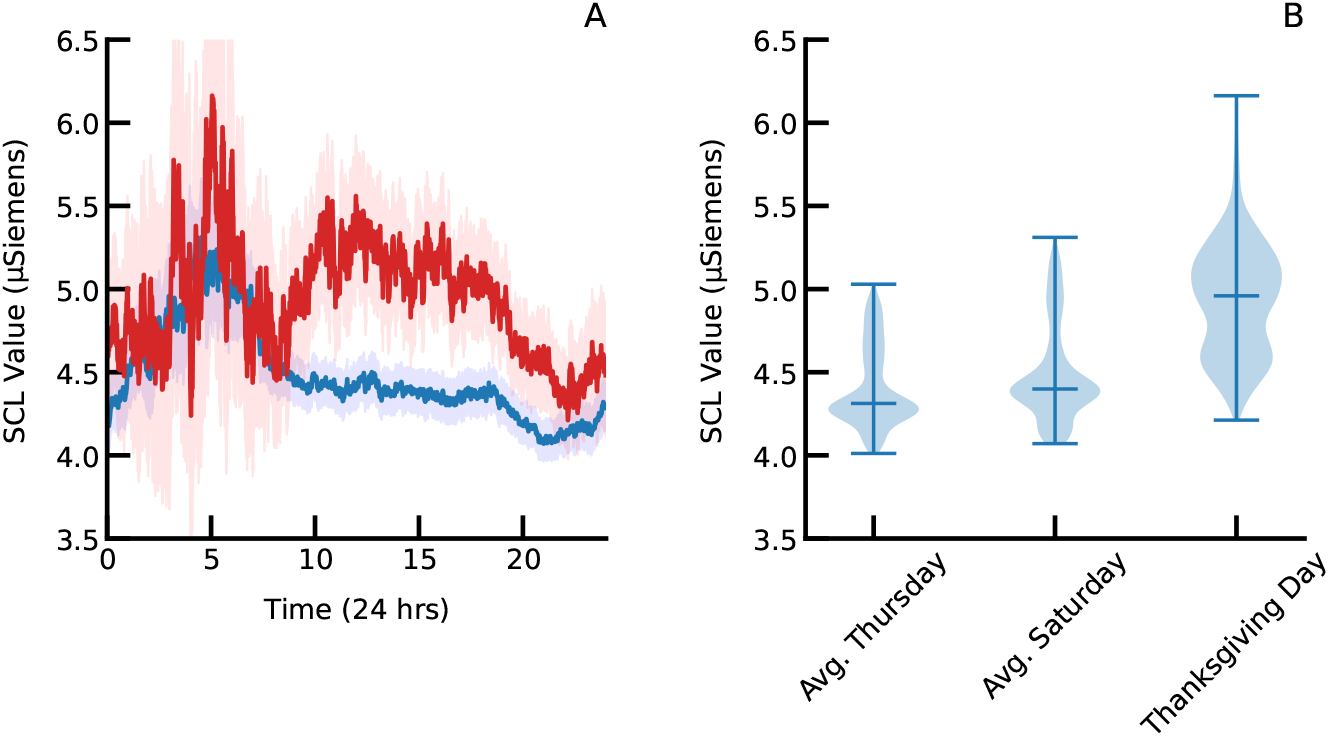
(A) The population mean SCL on Thanksgiving Day (red) and all other Thursdays in November and December (blue). Shaded areas represent 95% confidence based on standard error. (B) Distributions of mean minutely SCL values on Thanksgiving Day, all other Thursdays in November and December and Saturdays in November and December.

#### Super Bowl LVII

To analyse the effects of the Super Bowl in 2023, we segmented SCL data from one hour before kickoff until approximately one hour after the Trophy Award ceremony. We identified individuals whose state was Kansas (N=78), Pennsylvania (N=286) and Texas (N=572). Kansas is associated with the winning team, the Kansas City Chiefs^5^ and Pennsylvania is home to the defeated team, the Philadelphia Eagles. We select Texas as an example of a neutral state which is in the same timezone as Kansas. Mean SCL reveals elevation in SCL during the winning Field Goal, final whistle and trophy presentation for individuals with a home state of Kansas. A two-sided Mann-Whitney U test was performed to evaluate whether SCL differed between states, the results indicated that home state of Kansas had significantly greater SCL than Pennsylvania (z = [204,524], *p* = [.001]).

## Discussion

*In-situ* measurement of EDA confirms expected relationships between activity, environment, and events on SCL. These analyses reveal both diurnal patterns that are only explained in part by other sources of variance as well as predictably elevated responses to stressful or exciting life events including holidays, the Super Bowl, and “all-nighters”

As expected, body-mass index (BMI) was positively correlated with SCL, with those with higher BMI sweating more on average. This was the largest single effect in our multivariate model. Women had significantly higher SCL values than men; however, these differences were only pronounced in older people between the ages of 50-70 years. This suggests that the effect of menopause and hot-flashes are significant in a free-living population. Skin temperature was not significantly different in these age groups as both positive and negative changes in skin temperature [Molnar, 1975] may cancel out, whereas the same is not true for SCL.

The mean SCL was higher for weekends than weekdays (coeff = 0.007, *p*<0.001). However, this difference was driven primarily by higher SCL during the middle of the day with the largest effect from 12pm-6pm (coeff = -0.132, *p*<0.001) compared to 6am-12pm or 6pm-12am. A higher number of steps and skin temperature during these hours suggest alternative patterns of physical and outdoor activity at the weekends that could be driving the SCL differences. However, it is also possible that other factors, such as increased social interaction, decreased predictability in schedules, and less time in heating, ventilation, and air conditioned (HVAC) environments, may also contribute to the higher mean SCL on weekends.

As observed in prior work [Vieluf et al., 2021] we find a zenith in SCL during nighttime hours. This is despite the fact that the sensor was not logging SCL during minutes the subjects were detected as asleep (to save on battery power). It is known that elevated SCL can occur during certain sleep stages [Sano et al., 2014] (i.e., in non-REM slow-wave and non-REM stage 2 sleep); however, our results suggest that SCL is elevated during nighttime hours even when people are not sleeping. Furthermore, diurnal patterns of individuals who did not sleep at all show elevated SCL throughout the entire night. This result suggests that staying up late and/or missing nights of sleep is associated with sympathetic body responses.

Skin temperature was the most strongly related physiological or behavioral signal in our model to SCL; however, our recordings do show that SCL diurnal rhythms are not explained by the effects of thermoregulation, a result that is consistent with prior research [Vieluf et al., 2021]. Heart rate and HRV had only slight correlation with SCL, but the signs of those effects were in the expected directions with lower HRV correlated with higher SCL. Physical activity, in terms of steps was more substantially related, and physical activity was likely the cause of the maxima between the early morning and later in the day. These effects led to a two-bumped SCL rhythm best explained by a cosine with maxima at 6am and 6pm.

Examining holidays and sporting events such as Thanksgiving Day and the Super Bowl revealed expected elevations in SCL. To the authors’ knowledge, this is the first reporting of population-scale passive measurement of sympathetic activation using EDA. For Thanksgiving Day (a Federal holiday on the fourth Thursday in November), mean SCL was significantly higher than other Thursdays (*p* < 0.001) and Saturdays (p < 0.001) in November and December (Fig. 5). In addition to the social nature of Thanksgiving, this holiday is known for celebrating food and drink in abundance. Eating food increases parasympathetic activity whereas drinking alcohol reduces parasympathetic activity and increases sympathetic activity [Kloner, 2004, Olsson et al., 2021, O’Hara et al., 2020].

Turning focus to time-specific stressors, watching the Super Bowl, we see that while SCL on average throughout the game was not significantly higher for people from different states (see Fig. 6), the maxima in SCL were precisely time aligned with moments of expected high arousal, including the game winning field goal and trophy presentation to the winning team. It is also notable that people from both Kansas and Pennsylvania showed larger variations in SCL throughout the game than people from Texas (a neutral state). These results show that *in-situ* measurement of SCL shows meaningful differences at precise moments in time on a single day and not just when averaged throughout a longer time period.

**Figure 6:**
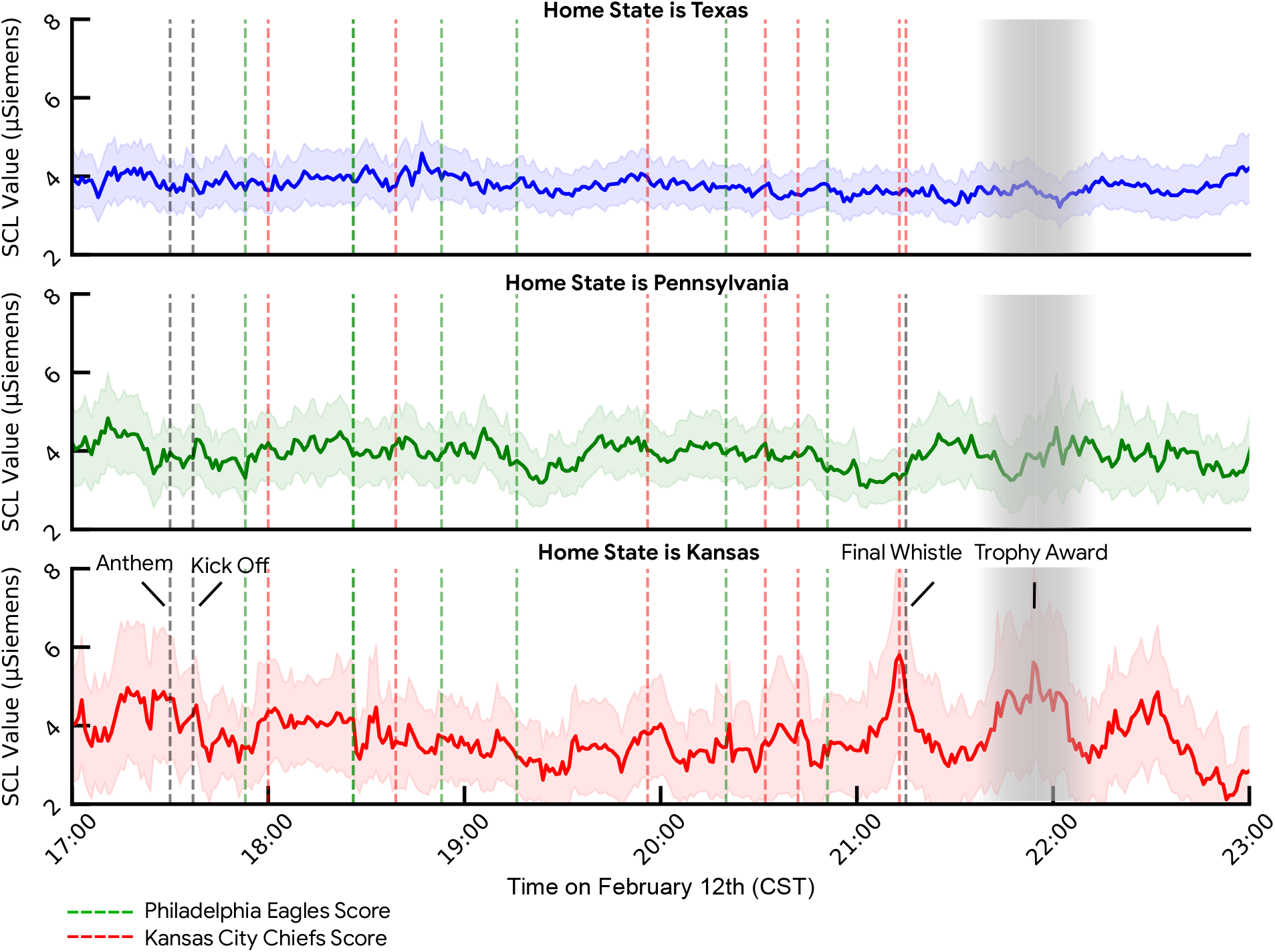
Mean SCL value for subjects whose home state was Texas, Pennsylvania (state of defeated team, the Philadelphia Eagles) and Kansas (state of the victorious team, the Kansas City Chiefs). SCL for those from Kansas was significantly higher during the final field goal, end of the game and trophy award than for those from Pennsylvania. Shaded areas represent 95% confidence based on standard error.

The present study does have several limitations. First, the data were collected from participants in the United States between October 1st 2022 to July 1st 2023. It is possible that SCL values may vary across seasons due to differences in the ambient conditions, and future studies should consider participants from other geographies. Secondly, the participants in this study were all users of the Fitbit Sense 2 devices which may represent a unique demographic of the general population. For example, Fitbit Sense 2 Users may be more active or health-conscious. As described in the Sensor Design section, the EDA sensor was turned off during periods in which. the device was in sleep mode. Therefore, the diurnal patterns need to be viewed with that caveat.

To our knowledge, these are the first large-scale observational profiles of EDA collected *in-situ*. Our results validate that dorsal wrist measured SCL is sensitive to sympathetic arousal in response to stressors, in addition to thermoregulation and sweating in response to physical activity. Our findings highlight the utility of using population-scale EDA sensing for observing *in situ* physiological responses to high arousal moments and in real-world settings. Overall, passive and continuous measurement of EDA via a wearable device appears a useful way for wearable devices to meaningfully impact both individual and population health. The ability to continuously quantify electrodermal activity introduces opportunities to scale physiological measures that are relevant to stress and stress pathology from laboratory to be translated to real world contexts.

## Data Availability

We support open science principles, but also recognize that it is often challenging to ensure participant privacy while also making data broadly available to the academic research community. We had to balance these considerations with the privacy of the participants and protection of their health data. Furthermore, although the data could be de-identified, some of the data streams could not be fully anonymized. We recognize that this is a limitation, but we felt that we needed to give the participants strong reassurance that their data would not be used for any other purpose than for the research at hand.

## Supplementary materials

### Methods

#### Sensor Design

Analyses used large scale continuous EDA data captured on the commercially available smartwatch, Fitbit Sense 2 which is enabled with a dorsal wrist-facing EDA sensor. This sensor is positioned on the underside of the device wherein it is designed to directly interface with the skin and allow a continuous measurement of the dorsal wrist impedance. This impedance signal is acquired at 200 samples-per-second, which is then passed through a boxcar filter and down-sampled to 25 samples-per-second. This signal is then shifted and scaled using a benchtop calibration mapping that compares measured known impedance values to the internal electrical impedances of the device, from which admittance is calculated. For the purposes of this study, the sensor data pulled from device for analysis was median-filtered minutely admittance values expressed in Siemens.

In addition to EDA, the Fitbit Sense 2 is equipped with multiple sensing modalities including: multi-path optical heart rate tracker, ambient light sensor, gyroscope, 3-axis accelerometer, on-wrist skin temperature sensor, and red and infrared sensors for oxygen saturation monitoring. These sensors are primary inputs into health and wellness features for this commercial device (such as heart rate or steps), and were included in our analysis. The nature of device operation (i.e. switching between different sensing modes) required that the EDA sensor be on during wake periods and off during sleep. Therefore, sleep EDA was excluded in our analysis.

The EDA sensors is used in the Fitbit Body Response feature which is not running during sleep. As a result the EDA sensors was only switched on during wake periods and off during sleep. Therefore, in our analyses we do not study EDA during sleep specifically and any measurements during “night time” hours was specifically for subjects who were awake during that time.

#### Subjects

In this study we collected data from individuals using Fitbit Sense 2 devices. All participants agreed to allow their deidentified data to be used for research purposes, as outlined in the Fitbit Terms and Conditions applicable at the time of data collection. We extracted age, sex, body-mass index, steps, EDA, IMU, heart rate and heart rate variability data, demographic information (age, gender) and device metadata (timezone, time). Body mass index (BMI) was calculated based on participants’ self-reported weight and height.

To study diurnal patterns we study a population of 15,349 US-based Sense 2 wearers (Men: 5,300, Women: 10,049; Mean Age: 49.8 years, Min: 18 years, Max: 91 years). The Fitbit Sense 2 launched in August 2022 and therefore we analyzed data from October 2022 to July 2023. These individuals had at least 4 hours of daytime wear on at least 90 days within this period. The age and BMI distributions of our sample broadly conform to the demographic profiles of the US general population (see Fig. 1). However, the sample did demonstrate a larger representation of women than men then is reflected in the general population (percent women: 64.4%; population distribution of women: 50.4%). The location (state) distribution also broadly reflects the population density within the US. Physical activity is known to influence EDA [Poh et al., 2010].

For analysis of the Super Bowl, we analyzed a subset of the 15,349 US-based Sense 2 wearers who were wearing their Sense 2 between 5pm and 11pm Eastern Standard time on February 12th 2023. Of these users there were 78 subjects from Kansas (men = 26, women = 52, mean age = 49.8 years, mean weight = 86.2kg), 286 from Pennsylvania (men = 104, women = 182, mean age = 51.7 years, mean weight = 86.3kg) and 572 from Texas (men = 209, women = 363, mean age = 50.3 years, mean weight = 86.4kg).

### Trier Social Stress Test Experiment

The Trier Social Stress Test (TSST) is considered a “gold-standard” for laboratory induced stress procedures. The protocol involved subjects being asked to participate in two stress inducing tasks: a mock job interview and a math task. The Trier Social Stress Test was conducted remotely during 2021. A remote version of the TSST had previously been validated [Eagle et al., 2021] and due to constraints imposed during the COVID pandemic, we used the remote protocol for this data collection.

The study protocol was approved by the Advarra Institutional Review Board (IRB No. Pro00054030). Informed consent was collected from all participants before the study and all subjects were compensated. Forty-five healthy adult English speakers (n=20 male) and were well-balanced across sex (males: M[SD]=39.6[9.2] years; females: M[SD]=37.0[8.5] years). Participants were excluded if they had any self-reported psychiatric conditions.

Two 20-minute interviewer-panel videos were created (one all male panel, one all female panel). The three-member interview-panel maintained a flat affect and moved minimally during the entire 20 minute videos. For each subject, the opposite-sexed video was used during the instructions, mock-job interview and arithmetic task portions of the study.

Subjects were instructed to wear a Fitbit Sense 2 for the hour prior to, and during, the study. Subjects joined a video-conferencing call with the experimenter, after which they were told to relax calmly at their desks for a 15 minute baseline period. Following this was a 10-minute anticipatory period consisting of five minutes of instructions for the mock job-interview and 5 minutes of interview preparation or note taking. Prior to administering the instructions, subjects were asked to join a break-out room (“study room”) where a second video-conferencing account was playing the opposite-sexed interviewer-panel video as the background, simulating a real interview-panel over video. The experimenter began recording the video call from this point onwards, and read the instructions to the subject from a different account. After the interview preparation period, the subject’s notes document was deleted and the subject was asked to enter the “study room” for the 10-minute stressor period consisting of a 5-minute mock-job interview and subsequent surprise 5-minute arithmetic task. During the stressor period, the experimenter’s audio and video feeds were turned-off and they communicated with the subject via chat messages. During the mock-interview, if the subject was quiet for 20 seconds then the experimenter sent a message asking them to please continue. After the arithmetic task was complete, the subject was asked to return to the “safe room” where they were debriefed for 10-minutes. Subjects were given a debriefing document with an explanation of the purpose of the study. Finally, subjects were asked to rest calmly again for a 15 minute recovery period.

### EDA Distributions

Fig. 7 shows the distribution (and Q-Q plot) for six-hourly SCL levels after a Box-Cox transformation. These data follow a normal distribution, as the null-hypothesis that they are not normally distributed can be rejected based on a two-sided Jarque-Bera chi^2^ test.

**Figure 7:**
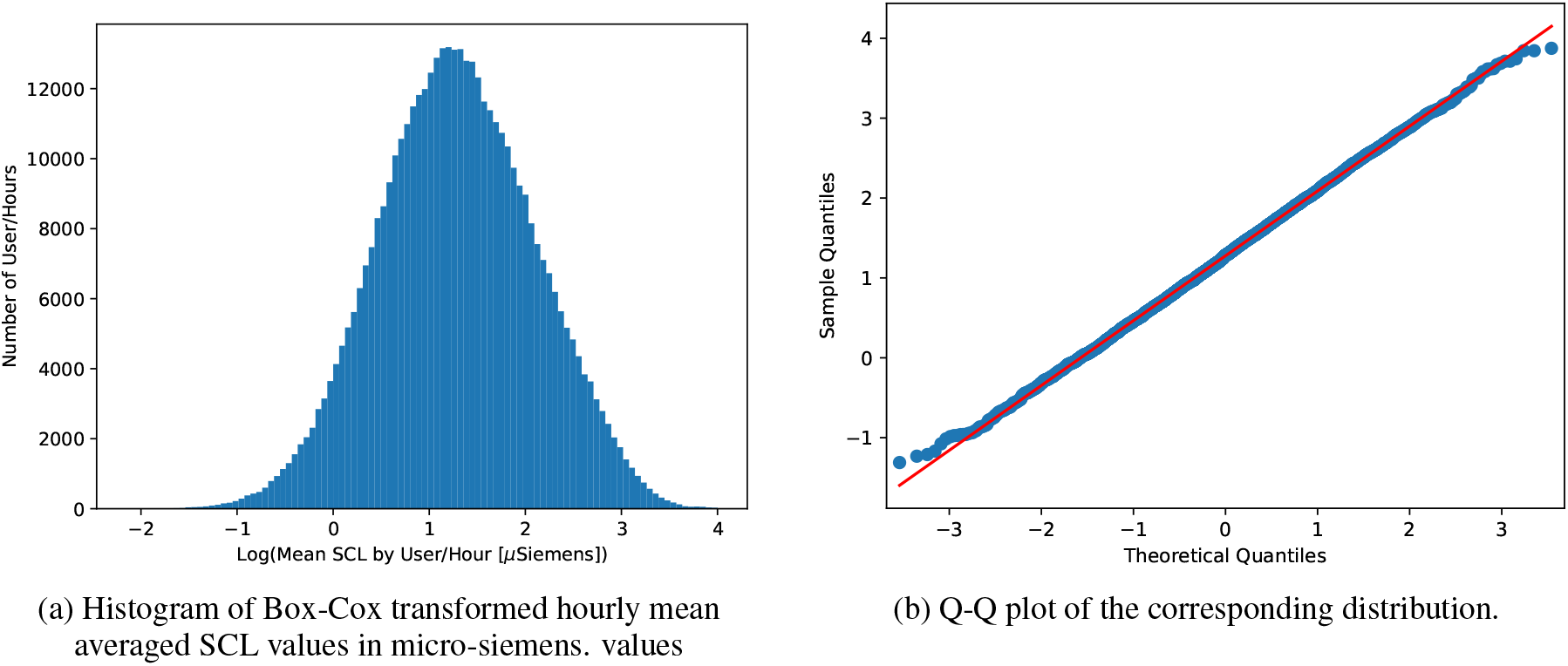
Distribution and Q-Q plots for six-hourly SCL values.

## Footnotes

1 https://affect.media.mit.edu/projectpages/iCalm/iCalm-2-Q.html

2 https://www.empatica.com/research/e4/

3 https://www.fitbit.com/global/us/products/smartwatches/sense

4 https://www.happyring.com/

5 Note: Kansas City is split between the states of Kansas and Missouri, but in this analysis we use Kansas.

